# Subtypes of functional brain connectivity as early markers of neurodegeneration in Alzheimer’s disease

**DOI:** 10.1101/195164

**Authors:** Pierre Orban, Angela Tam, Sebastian Urchs, Melissa Savard, Cécile Madjar, AmanPreet Badhwar, Christian Dansereau, Jacob Vogel, Amir Schmuel, Alain Dagher, Sylvia Villeneuve, Judes Poirier, Pedro Rosa-Neto, John Breitner, Pierre Bellec, for the Alzheimer’s Disease Neuroimaging Initiative, and the Pre-symptomatic Evaluation of Novel or Experimental Treatments for Alzheimer’s Disease Program

**Author notes:** Data used in preparation of this article were obtained from the Alzheimer’s Disease Neuroimaging Initiative (ADNI) database (adni.loni.usc.edu). As such, the investigators within the ADNI contributed to the design and implementation of ADNI and/or provided data but did not participate in analysis or writing of this report. A complete listing of ADNI investigators can be found at: http://adni.loni.usc.edu/wpcontent/uploads/how_to_apply/ADNI_Acknowledgement_List.pdf. Data used in preparation of this article were obtained from the Pre-symptomatic Evaluation of Novel or Experimental Treatments for Alzheimer’s Disease (PREVENTAD) program (http://www.prevent-alzheimer.ca), data release 2.0 (November 30, 2015). As such, the investigators of the PREVENT-AD program contributed to the design and implementation of PREVENT-AD and/or provided data but did not participate in analysis or writing of this report. A complete listing of PREVENT-AD Research Group can be found in: https://preventad.loris.ca/acknowledgements/acknowledgements.php?date=[2015-11-30].

## Abstract

**Highlights:**

- Reliable functional brain network subtypes accompany cognitive impairment in AD
- Symptom-related subtypes exist in the default-mode, limbic and salience networks
- A limbic subtype is associated with a familial risk of AD in healthy older adults
- Limbic subtypes also associate with beta amyloid deposition and ApoE4

**In Brief:** We found reliable subtypes of functional brain connectivity networks in older adults, associated with AD-related clinical symptoms in patients as well as several AD risk factors/biomarkers in asymptomatic individuals.

**Summary:** The heterogeneity of brain degeneration has not been investigated yet for functional brain network connectivity, a promising biomarker of Alzheimer’s disease. We coupled cluster analysis with resting-state functional magnetic resonance imaging to discover connectivity subtypes in healthy older adults and patients with cognitive disorders related to Alzheimer’s disease, noting associations between subtypes and cognitive symptoms in the default-mode, limbic and salience networks. In an independent asymptomatic cohort with a family history of Alzheimer’s dementia, the connectivity subtypes had good test-retest reliability across all tested networks. We found that a limbic subtype was overrepresented in these individuals, which was previously associated with symptoms. Other limbic subtypes showed associations with cerebrospinal fluid Aβ_1-42_ levels and ApoE4 genotype. Our results demonstrate the existence of reliable subtypes of functional brain networks in older adults and support future investigations in limbic connectivity subtypes as early biomarkers of Alzheimer’s degeneration.

## Introduction

Alzheimer’s disease (AD) is a chronic neurodegenerative condition that gives rise to the most common form of dementia, with severe memory and cognitive impairments. Importantly, the clinical expression of AD becomes apparent only decades after the development of neuropathological processes, such as the accumulation of amyloid-β (Aβ) plaques and tau neurofibrillary tangles. The long preclinical buildup of AD pathology presents an opportunity to prevent, rather than repair, neurodegeneration (Dubois et al., 2016; Sperling et al., 2012). Functional brain connectivity measured with resting-state functional magnetic resonance imaging (rs-fMRI) may capture early synaptic dysfunction in AD (Selkoe, 2002; Tampellini, 2015) and is thus a promising candidate biomarker for AD (Badhwar et al., 2017; Brier et al., 2014; Jones et al., 2016; Vemuri et al., 2012). However, the current literature has largely relied on comparisons between group averages of patients and cognitively healthy individuals. Such cross-sectional analyses neglect the considerable phenotypic heterogeneity present both in patient and control populations. The primary objective of this work was to characterize the heterogeneity of functional brain connectivity in older adults, and identify network subtypes associated with AD at the clinical and preclinical stages.

A prevalent model of AD postulates that symptoms arise as a consequence of disruptions in distributed networks, rather than local, circumscribed alteration in neural processing (Delbeuck et al., 2003; Seeley et al., 2009). The seminal work of (Greicius et al., 2004) in symptomatic AD demonstrated alterations in functional brain connectivity in the so-called default-mode network (DMN), whose topography overlaps substantially with patterns of end-stage Aβ deposition (Buckner et al., 2005). A recent meta-analysis concluded to convergent evidence across over 30 publications looking at functional brain connectivity in clinical cohorts, i.e. patients with mild cognitive impairment or AD dementia, and confirmed the DMN as a key affected brain component (Badhwar et al., 2017). Connectivity disturbances in other large-scale brain networks were also found consistently, in particular in the limbic and salience networks. At a preclinical stage, rs-fMRI connectivity has been shown to be impacted in cognitively healthy older adults at risk of AD due to abnormal levels of cerebrospinal fluid (CSF) Aβ_1-42_ or tau proteins (Jiang et al., 2016; Wang et al., 2013), increased cerebral Aβ deposits (Elman et al., 2016), and presence of apolipoprotein E ε4 allele - ApoE4 (Sheline et al., 2010), the major genetic risk factor in sporadic AD. A familial history of sporadic AD in first-degree relatives is the second most important risk factor of AD (Tanzi, 2012), and was shown to impact DMN connectivity even in ApoE4 non-carriers, thus highlighting additional genetic risk factors (Wang et al., 2012).

Despite mounting evidence of rs-fMRI as an early marker of AD, the current literature neglects the considerable heterogeneity present in both patients and controls. Post-mortem histological examination of AD pathology in brain tissue samples (Hyman et al., 2012) indeed does not align closely with clinical diagnosis. Between 30 and 50% of patients diagnosed with AD dementia in fact do not present Alzheimer’s pathology, depending on the level of neuropathological confidence (Beach et al., 2012).

Conversely, the same study reported that close to 40% of patients diagnosed with non-AD dementia show minimal signs of AD pathology. Some cognitively healthy persons included in control groups may also suffer from preclinical AD, with 10% to 30% of them having Aβ deposition in their brain (Chételat et al., 2013), and some of them exhibiting high loads of neurofibrillary tangles (Mufson et al., 2016). Data-driven analysis of structural MRI subtypes in AD further showed that symptomatic heterogeneity (Belleville et al. 2007; Scheltens et al. 2016) is at least partly related to different modes of atrophy spreading in AD (Dong et al., 2017; Zhang et al., 2016). Complementary to subtypes of brain atrophy, a recent work (Doan et al., 2017) showed that connectivity subtypes can also be observed using diffusion magnetic resonance imaging, in patients suffering from AD dementia, MCI or subjective cognitive impairment, with subtypes accompanying the severity of cognitive impairment. The established heterogeneity in structural brain degeneration calls to re-examine the current evidence for rs-fMRI as an AD biomarker using a subtyping approach.

The overarching goal of the present work was to identify one or multiple subtypes of functional brain connectivity associated with AD, either at a clinical or preclinical stage. We first applied a data-driven cluster analysis to identify subgroups of subjects with homogeneous subtypes of brain connectivity within a mixed cohort of 130 subjects, including patients with AD dementia (AD subjects, N = 21), patients with mild cognitive impairment (MCI subjects, N= 44), and elderly healthy controls (HC subjects, N= 65) (Figure 1, Table 1). This mixed cohort, referred to as the ADNI2-MTL sample, poolsdatafrom 2 sites of ADNI2 (ADNI2a and ADNI2b) and 3 studies conducted at Montreal sites (MNI, CRIUGMa and CRIUGMb), in an attempt to extract robust subtypes that will generalize well to new studies (Orban et al., 2017). For each brain network and connectivity subtype, we tested whether a particular subtype was associated with the presence of mild or severe symptoms. We then investigated if the subtype membership was a reliable quantity using test-retest data in an independent sample of 231 cognitively healthy older adults, with a familial history of AD (FH subjects) (Orban et al., 2015). As those subjects are at risk for AD, we tested if the subtypes associated with symptoms would already be overrepresented in the asymptomatic FH cohort. We further tested in these FH subjects the association between functional subtypes and known biomarkers/risk factors of AD, namely CSF Aβ_1-42_, Tau levels as well as ApoE4 genotype.

**Figure 1.**
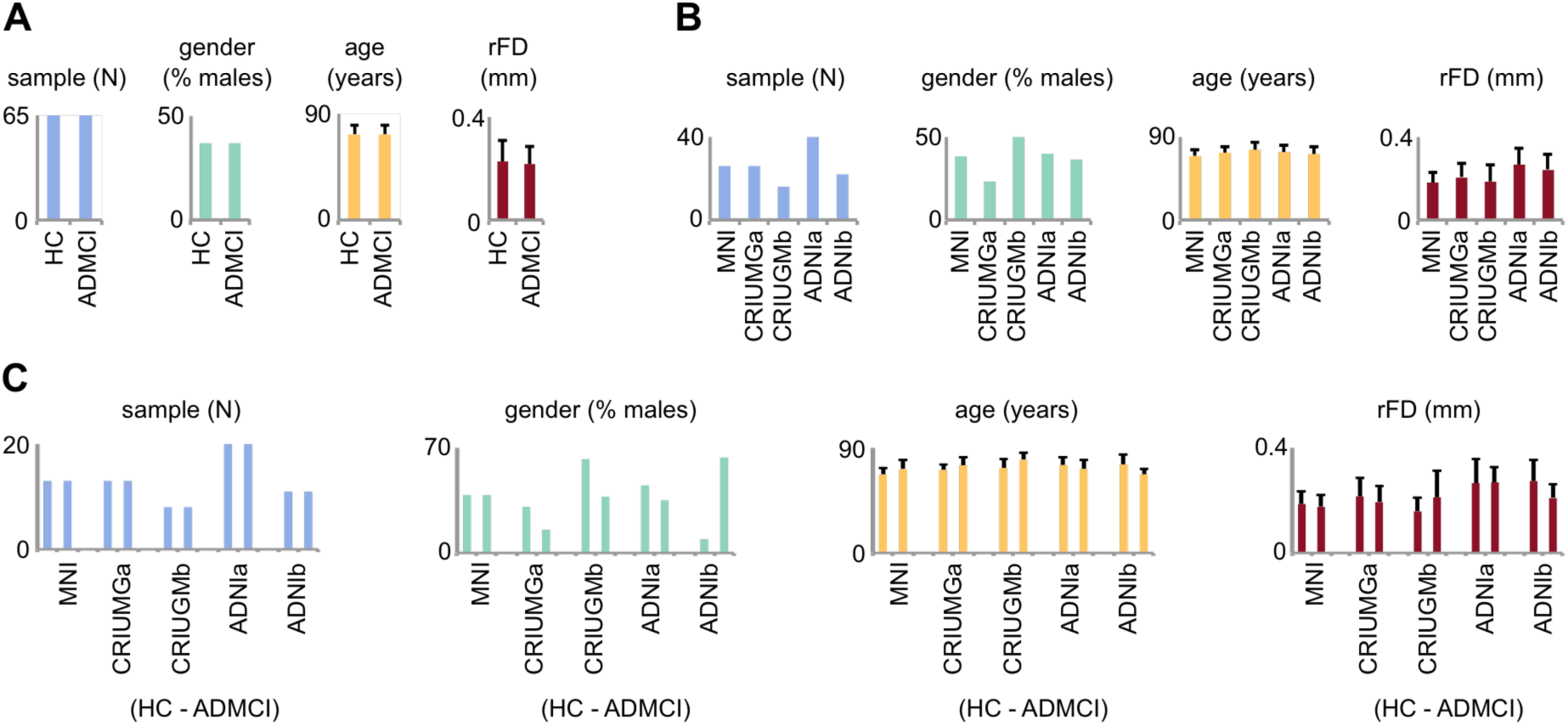
Matching between ADMCI patients and HC. (A) Patients and controls were matched with respect to sample size, gender, age and motion levels after scrubbing (residual frame displacement, rFD). (B) Between-site differences on4 such variables are shown irrespective of clinical status. (C) The number of patients and controls are perfectly balanced within sites.

**Table 1.**
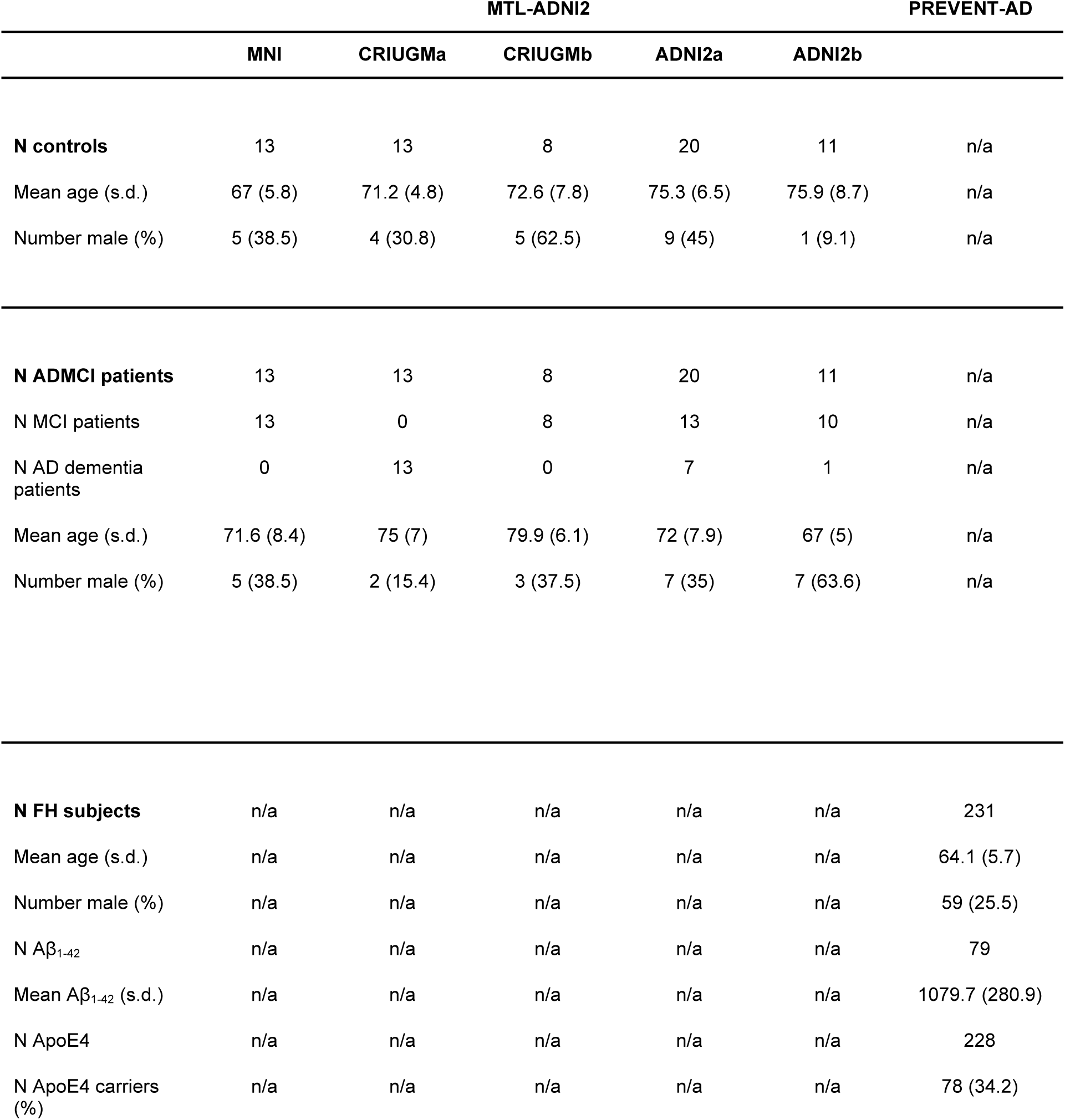
Demographics. Basic demographics (sample size, mean age, sex proportions) are given for the HC, ADMCI and FH groups. Levels of CSF Aβ_1-42_ and proportions of ApoE4 carriers are given for FH subjects.

## Results

### Subtypes of functional brain networks

To identify subtypes of functional brain networks, we first generated individual functional connectivity maps for seven large-scale networks together covering the entire brain (Figure 2A-B). These reference networks were obtained from an independent dataset from 200 healthy young subjects (Bellec et al. 2015), and were labeled as cerebellar, limbic, motor, visual, default-mode, fronto-parietal and salience networks. For each network, a hierarchical cluster analysis was applied on 130 individual network maps from the ADNI2-MTL dataset, after regression of phenotypic and site confounds, in order to identify subgroups of subjects with homogeneous brain maps. Visual inspection suggested the presence of at least three voxelwise connectivity subgroups (Figure 2C-D). A brain map averaged across all subjects within a subgroup defined a subtype of network connectivity, highlighting specific brain areas that differed between that subgroup and the overall population average (Figure 2E). Subtype maps revealed high connectivity with their reference network, yet also exhibited noticeable variations. These differences were not only observed in the associated network (within-network connectivity) but also in other brain areas (between-network connectivity). For instance, subtypes of the DMN could be distinguished from one another not only in terms of connectivity levels within the precuneus or anterior medial prefrontal cortex, two key nodes of the default-mode network, but also with regards to connectivity strength in the anterior cingulate, associated with the salience network. For each network, we generated the spatial correlations between individual connectivity maps and each average subtype map, hereafter referred to as weights (Figure 2F). These continuous subtype weights revealed that some individual maps were highly correlated with the subtypes, while others had only milder correlations, sometimes of similar amplitude for different subtypes. The subtype decomposition was therefore a discrete approximation of a continuous distribution of individual maps, rather than a set of clear-cut entities.

**Figure 2.**
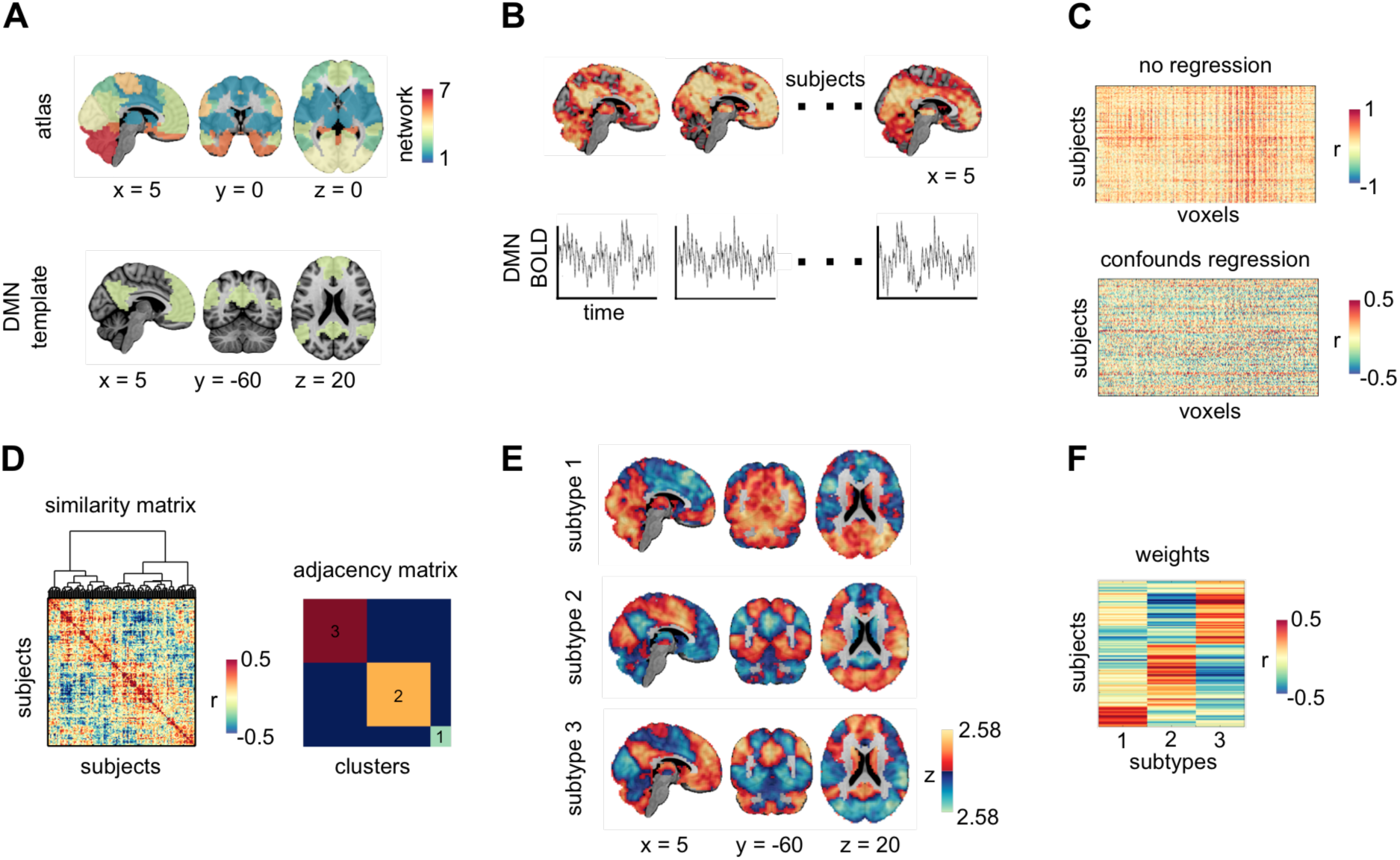
Extraction of subtypes and weights. (A) Functional subtypes were identified separately for 7 networks delineated at the whole-brain level in an independent sample of healthy subjects. The procedure is shown for the default-mode network (DMN). (B) Network-based connectivity maps were computed for each subject through the correlation of every voxel’s time course of activity with the average signal in the reference network. (C) Site, gender, age and motion were regressed out from functional connectivity maps across subjects. (D) A hierarchical cluster analysis was conducted to identify 3 homogeneous subgroups of subjects with similar connectivity maps. (E) Difference subtypes show how the average connectivity maps of each separate subgroup of subjects differ from the grand average. (F) Weights consisted in correlations between the connectivity maps of every subject with that of each subtype.

A comparison of clustering outcomes for the seven networks revealed that 3 subgroups of subjects at least could be evidenced in all networks (Figure 3A). As observed for the DMN, subtype maps showed distributed variations inside and outside the network of reference for all networks. While between-subject correlation values had similar amplitudes across networks, the size of the subgroups varied from one network to another. We tested the correspondence of subject clustering solutions between networks by computing the adjusted rand index (ARI) for all pairwise comparisons (Figure 3B). The near-chance level of this metric (0.04 ± 0.04) demonstrated that subjects with similar connectivity maps for a given network did not have particularly similar maps for other networks, thus highlighting heterogeneity in functional brain connectivity patterns.

**Figure 3.**
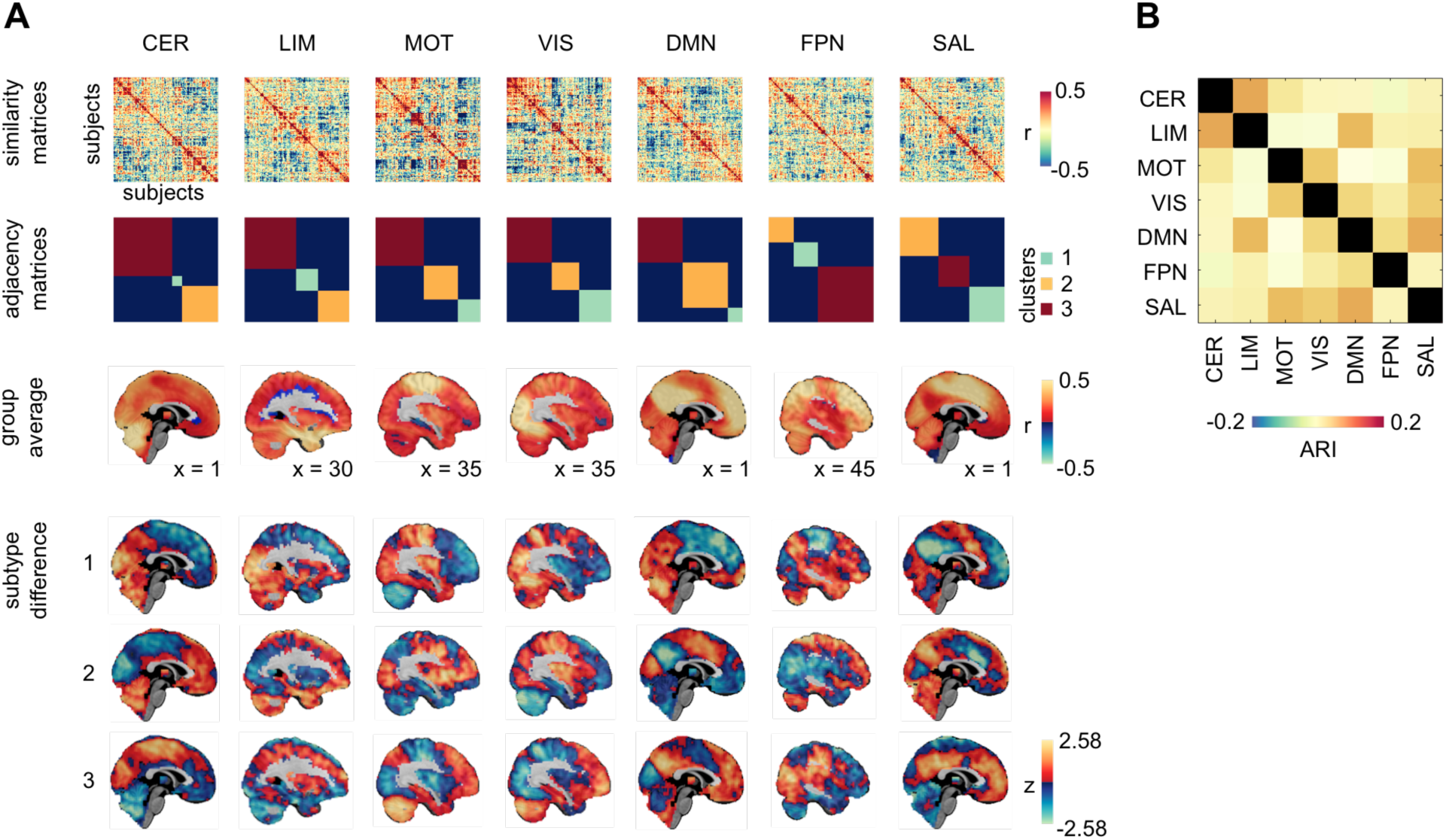
Correspondence of cluster (subtype) solutions across networks. (A) For each of the 7 networks (columns) are given the similarity matrix that shows the similarity of network connectivity maps between all pairs of subjects (first row), the adjacency matrix that reveals homogeneous subgroups of subjects identified by cluster analysis (second row), the average network connectivity map for all subjects (third row), and the difference subtype connectivity maps obtained by differences between the group average and the average connectivity maps for each subgroup of subjects (fourth to sixth rows). (B) The adjusted rand index (ARI) reveals the correspondence of subject clustering solutions between all pairs of networks. CER, cerebellum; LIM, limbic; MOT, motor; VIS, visual; DMN, default-mode; FPN, fronto-parietal; SAL, salience.

### Brain network subtypes are associated with clinical symptoms

Given the observation that subtypes reflected both continuous and discrete phenomena, we adopted a dual statistical evaluation of their association with clinical symptoms in ADMCI subjects (Figure 4). In the former case, differences in average subtype weights between ADMCI and HC were assessed independently for each subtype of the seven reference networks, using a linear regression model. Significant associations were found for one limbic, two default-mode and two salience subtypes (q < 0.05 with FDR correction over 21 network subtypes), in line with our expectations. An uncorrected effect was also seen for an additional limbic subtype (p < 0.05). Effects were of medium size (0.09 < Cohen’s f2 < 0.25). Of these six subtypes, half of the associations with symptoms were positive (i.e. higher average weight load in ADMCI persons) and the remainder negative (i.e. lower average weight load in ADMCI patients). Instances of positive and negative associations with symptoms were observed in all three aforementioned networks.

**Figure 4.**
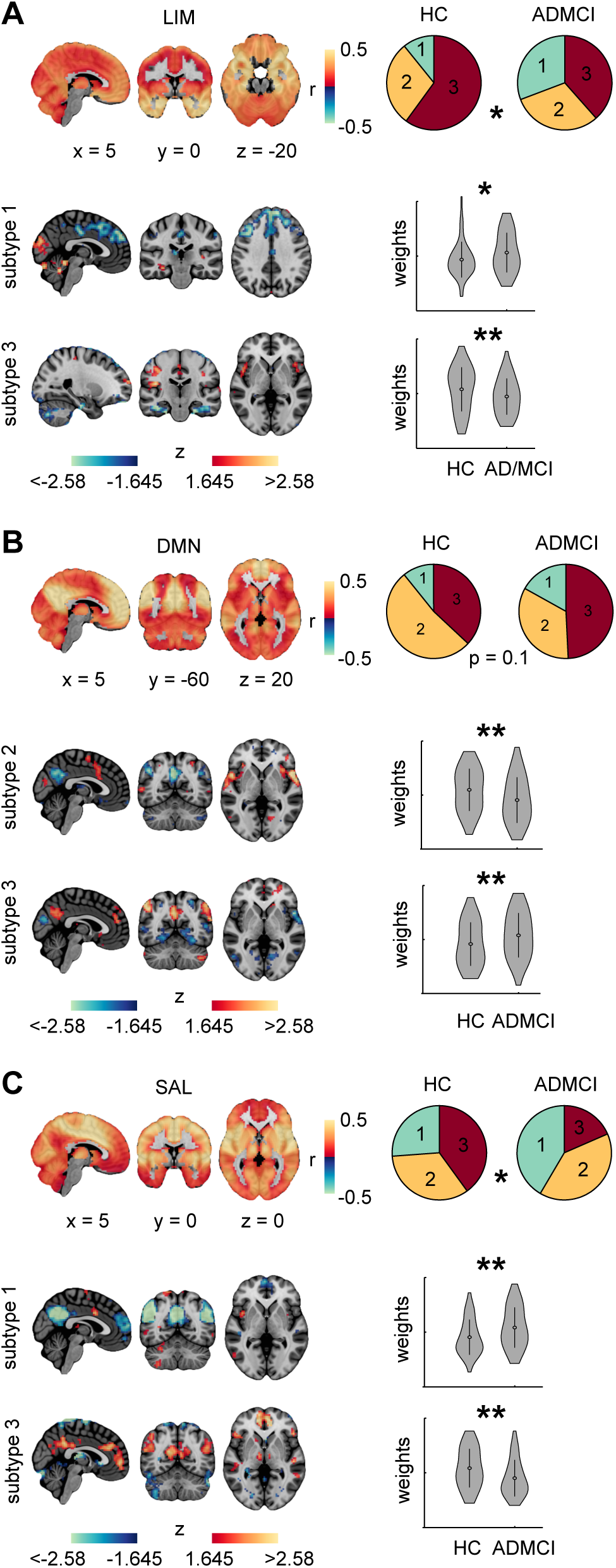
Functional network subtypes associated with clinical symptoms. Significant associations with ADMCI were found in the limbic (A), default-mode (B) and salience (C) networks. For each network are shown the group average connectivity map and the connectivity subtypes that are significantly more or less present in ADMCI patients than controls (difference maps are given). Pie charts report the distributions of subjects across subtypes in each group. Violin plots show the distribution of weights in the two groups for each subtype with a significant association. ** and * respectively denote significance at qFDR<0.05 and p<0.05 (uncorrected).

A general observation was that subtypes positively associated with symptoms (PAS) had increased within-network connectivity but decreased between-network connectivity as compared to sample averages of networks. The PAS limbic subtype was notably defined by increased hippocampal connectivity (within-network) but decreased connectivity in dorsomedial prefrontal areas located in the DMN (between-network). An inverse pattern was seen in subtypes negatively associated with symptoms (NAS). The NAS limbic subtype had decreased connectivity in the hippocampus but increased connectivity in the insula. Subtypes of the default-mode and salience network provided mirror pictures of PAS and NAS connectivity profiles. Decreased connectivity in the posterior cingulate and medial prefrontal region relative to the sample average was NAS for the default-mode network but PAS for the salience network. Similarly, decreased connectivity in the insula and anterior cingulate cortex compared to the sample average was evidenced to be NAS for the salience network but PAS for the default-mode network.

Statistics on discrete effects provided concordant effects at uncorrected thresholds. For each network, we evaluated with Chi2 tests whether ADMCI and HC subjects were distributed unevenly across subtypes. Unequal distributions were seen for the limbic (p < 0.05), default-mode (p = 0.1) and salience (p < 0.05) networks. Effect sizes were in the small-to-moderate range, with Cramer’s V values of 0.27, 0.19 and 0.24 in the limbic, default-mode and salience networks, respectively.

### Connectivity maps in FH subjects are reproducibly matched to subtypes from the clinical cohort

We assessed the reliability of matching connectivity maps in FH subjects from the PREVENT-AD cohort with the subtypes defined in the MTL-ADNI2. We thus generated individual functional connectivity maps separately for two runs, in each of the seven networks. Weights were computed for individual network maps, indicating their similarity with each of the 21 network subtypes previously defined in the MTL-ADNI2 sample (Figure 5). Intraclass correlations (ICC) indicated a fair-to-good correspondence of subtype weights between runs. Weights of all network subtypes had ICC values > 0.45 (max = 0.68, mean = 0.56), except for the PAS salience subtype (0.29). The default-mode and limbic PAS subtype weights had ICCs of 0.50 and 0.55, respectively.

**Figure 5.**
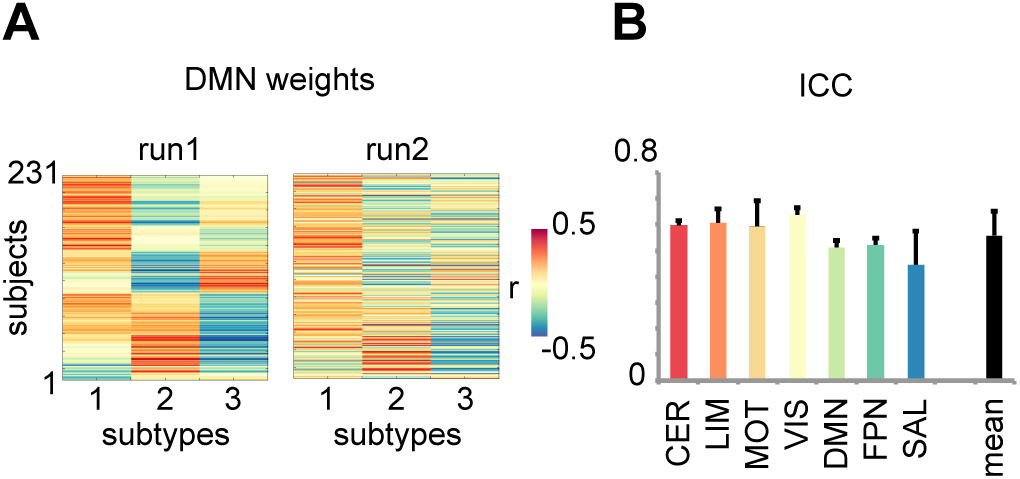
Reliability of subtype matching in FH subjects. (A) Matching of connectivity maps in FH subjects with subtypes found in the mixed population of ADMCI patients and controls is shown for the DMN in two separate runs. (B) Test-retest between runs was determined with intra-class correlation (ICC), showing fair-to-good correspondence across networks and subtypes.

### Subtypes are associated with biomarkers of AD in FH subjects

We next examined the possibility that cognitively healthy FH older adults already exhibited PAS subtypes, and more so than typical healthy elderly individuals. Individual functional connectivity maps were averaged for the two separate runs in 231 FH subjects from the PREVENT-AD cohort. For each of the three networks found to be associated with clinical symptoms, FH subjects were matched to network subtypes defined in the MTL-ADNI sample based on maximal weights. Distributions of FH subjects across subtypes were not significantly different than those of either ADMCI or HC participants in the default-mode and salience networks (Figure 6A). However, proportions of FH subjects across limbic subtypes differed significantly from those of typical HC older adults (q < 0.05) but not from ADMCI patients (p = 0.9).

**Figure 6.**
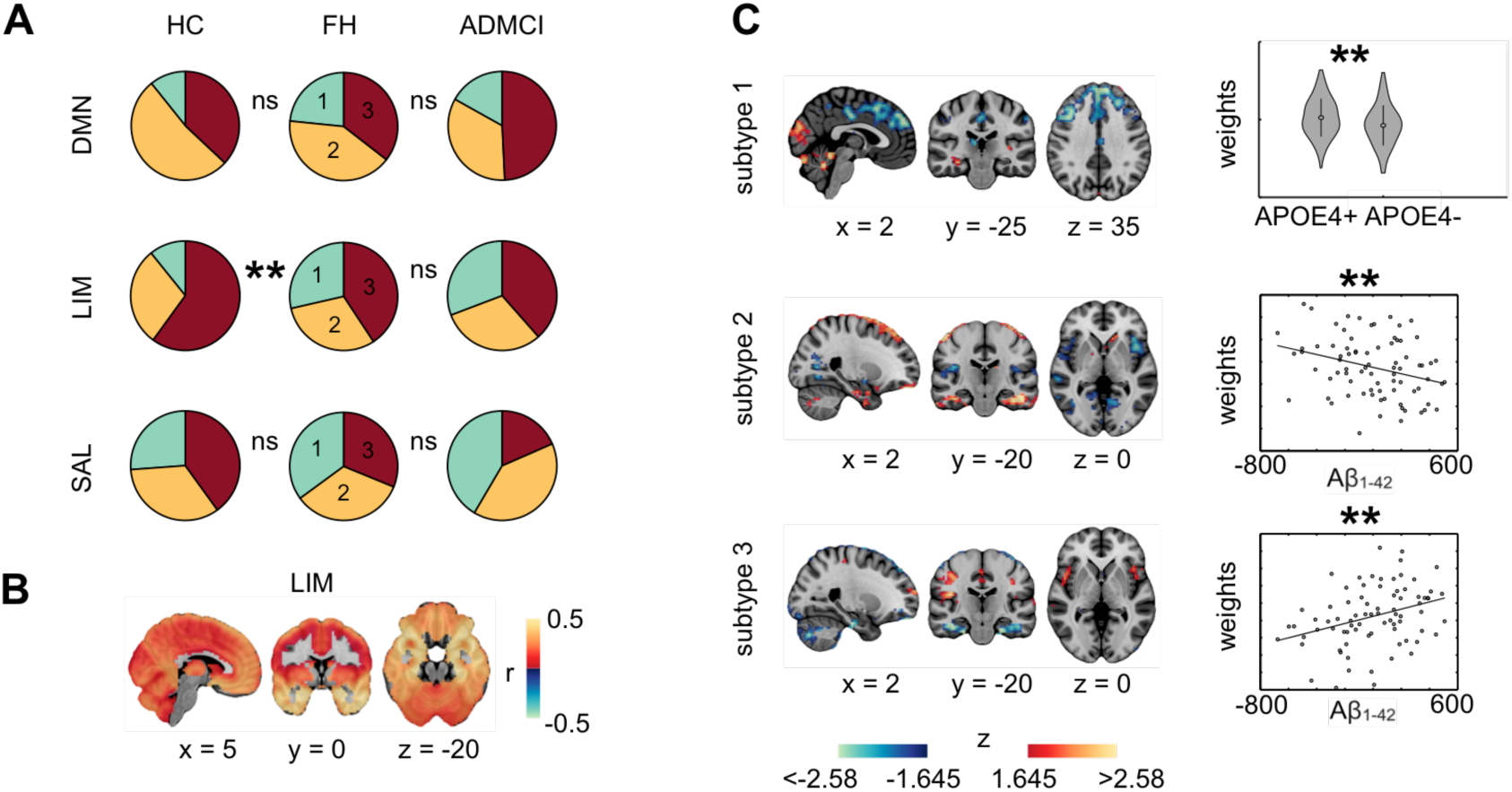
Connectivity subtypes in FH subjects. (A) Pie charts show that FH subjects differ from controls but not ADMCI patients in their distribution across subtypes for the limbic network (B). (C) Three distinct limbic network subtypes show either positive or negative associations with ApoE4 status or CSF Aβ_1-42_ levels. ** denotes significance at q^FDR^<0.05.

The idea that connectivity subtypes might reflect a covert pathological AD process in cognitively healthy elderly individuals would be reinforced by the observation that such connectivity profiles correlate with known biomarkers of AD. We thus further investigated the relationship between connectivity subtypes and APOE genotype (N = 228) as well as CSF levels of Aβ_1-42_, tTau and pTau (N = 59) (Figure 6C). Surprisingly, APOE allele 4 carriers showed less association than non carriers with the limbic PAS subtype (q < 0.05), with a small effect size (Cohen’s f2 = 0.04). However, findings consistent with predictions were observed for CSF Aß42 levels and another limbic subtype. Subjects with high levels of CSF Aβ_1-42_ had limbic connectivity maps that resembled more the NAS limbic network (q < 0.05; Cohen’s f2 = 0.13). Low levels of CSF Aβ_1-42_ were associated with another limbic subtype that shared some similarities with the PAS limbic subtype, for instance increased hippocampal connectivity (q < 0.05; Cohen’s f2 = 0.1). No associations were found between Tau or pTau CSF levels and any subtype of either the limbic, default-mode or salience networks.

## Discussion

### Capturing heterogeneity through subtyping

Our subtyping approach was motivated by the lack of specificity and sensitivity of a clinical diagnosis of AD dementia against a histopathological diagnosis of AD pathology (Beach et al., 2012) and the variability of cognitive and neurobiological alterations in AD (Lam et al., 2013; Scheltens et al., 2016). As done previously for structural atrophy patterns (Dong et al., 2017; Hwang et al., 2016; Zhang et al., 2016) and white matter structural dysconnectivity (Doan et al., 2017), we employed a subtype analysis that identified subgroups of subjects sharing similar functional brain connectivity, in a fully data-driven way and irrespective of clinical diagnosis. This is an important conceptual difference with more traditional cross-sectional comparisons between clinical cohorts, which assumes some homogeneity in connectivity within each group, e.g. (Badhwar et al., 2017, 2016; Jones et al., 2016; Korolev et al., 2016). Improved characterization of the inherent heterogeneity of brain dysconnectivity in AD will ultimately facilitate more personalized diagnosis and treatment. This new line of inquiry is made possible by large neuroimaging databases such as the ADNI, and will become increasingly important with the emergence of populational cohorts with associated neuroimaging repositories, such as the UK biobank (Miller et al., 2016).

### Association between connectivity subtypes and clinical symptoms

Using rs-fMRI, we identified functional brain connectivity subtypes associated both positively and negatively with symptoms. A variety of causal mechanisms may explain such associations, which may co-exist. An association may reflect the direct progression of AD neurodegeneration in the brain (Jones et al., 2016), the presence of comorbidities (Profenno et al., 2010), as well as some form of cognitive reserve, or lack thereof (Stern, 2006). The existence of an association in itself is not enough to disambiguate between these different interpretations. Associations between connectivity subtypes and symptoms were selectively detected in the default-mode, salience and limbic networks. These three networks have consistently been reported in the literature as altered in patients with AD dementia or MCI, see (Badhwar et al., 2017; Vemuri et al., 2012) for reviews. The associated subtype maps pointed at changes both within networks, e.g. higher intra-network connectivity in PAS DMN subtype, and between networks, e.g. decreased inter-network connectivity in PAS DMN subtypes with regions of the salience network.

### Translation of connectivity subtypes from clinical to non-clinical individuals

The distribution of connectivity subtypes in a group of cognitively normal FH individuals was found to resemble more the subtype distribution of a patient group than that of a control group. This observation was made only for the limbic network, but not the default-mode and salience networks. Assuming functional connectivity subtypes partly reflect the progression of AD pathology, finding early dysconnectivity in the limbic network is consistent with the Braak staging of neurodegeneration (Braak and Braak, 1991) and the increased risk of sporadic AD due to family history (Tanzi, 2012). Conversely, the limbic subtype negatively associated with symptoms was under-represented in FH individuals, and was shown to positively associate with CSF Aβ_1-42_ levels. Taken together, these associations support the notion that different subtypes of limbic connectivity reflect the progression of AD pathophysiology at a preclinical stage. A finding that was more difficult to interpret was that ApoE4 carriers had significantly less weight on the limbic subtype positively associated with symptoms. With previous literature on ApoE4 and resting-state connectivity sometimes reporting contradictory findings (Filippini et al., 2009; Sheline et al., 2010), we believe longitudinal data on a large cohort would be necessary to clarify the relationships among resting-state connectivity, Aβ deposition and ApoE4 status.

### Generalization of brain connectivity subtypes across datasets

The translation of connectivity across cohorts raises the question of generalization across scanning sites. Research has indeed indicated that multisite scanning generates substantial site-specific bias in connectivity measures (Dansereau et al., 2017; Yan et al., 2013). In our multisite clinical sample, we took great care to control for confounding site effects on brain connectivity subtypes. The identification of network subtype was thus invariant to scanning site to a large extent. However, the cohort of individuals at risk of AD due to their familial history was entirely scanned at a single site. The fact that we found associations with known biomarkers or risk factors of AD specifically in the limbic network supports that brain connectivity subtypes are fairly robust to site effects. Subtype weights also had good test-retest reliability in the PREVENT-AD cohort, although the subtype maps were generated on ADNI-MTL. Important areas for future work will be to identify imaging protocols that further minimize differences in brain connectivity subtypes across scanners.

### Finer subtypes

Groups of patients that defined subtypes did not overlap a lot across networks, including for subtypes positively associated with symptoms. There is thus some degree of independence between subtypes across networks, possibly reflecting heterogeneity of neurodegeneration across patients. Even though we estimated only 3 subtypes per network, there are still a very large number of possible combinations of subtypes across 7 networks. Subtype maps being an average of a subgroup of subjects, a minimum number of 20 subjects seems warranted to stabilize the subtype maps. The total sample size of our discovery dataset thus constrained the maximal number of subtypes we could feasibly investigate. We thus decided to use low numbers of subtypes and networks for this first evaluation of the feasibility of functional subtypes in AD, yet higher numbers could be explored in a larger sample.

### Multi-network and multimodal subtypes

A natural extension of this work would be to integrate subtypes across multiple networks, imaging modalities and measures into a single predictor of AD status. Associations with clinical symptoms or AD biomarkers reported here had weak to moderate effect sizes, despite reaching statistical significance. Recent state-of-the-art model of progression from MCI to dementia indeed merge biomarkers across multiple domains, including cognitive evaluations, imaging and plasma markers (Korolev et al., 2016). High-dimensional imaging biomarkers such as structural and diffusion MRI are amenable to subtyping (Doan et al., 2017; Hwang et al., 2016; Zhang et al., 2016). We believe that subtyping could be used in the near future to identify a highly accurate multimodal predictor of AD, both for diagnosis and prognosis purposes. Resting-fMRI will likely contribute to such a multimodal predictor, as it is uniquely sensitive to brain function, at least compared to other MRI modalities. Our findings suggest that limbic subtypes in particular are promising biomarkers for the purpose of early AD diagnosis.

### Conclusions

The present work demonstrates that rs-fMRI can be used to subtype the heterogeneity of functional networks in older adults. We found that subtypes have a good test-retest reliability and associate with symptoms in patients suffering from MCI or AD dementia. We also found that subtypes associate with various biomarkers and risk factors of AD in cognitively normal individuals: familial history of AD dementia, beta amyloid deposition, ApoE4 status. Our findings support the notion that rs-fMRI subtypes are sensitive to AD progression up to the preclinical stage, and may contribute to future efforts towards an accurate early diagnosis of AD using multimodal biomarkers.

## Experimental procedures

### Participants

The MTL-ADNI2 multisite sample aggregated data from 5 different studies: 3 samples from the Montreal area (one from the Montreal Neurological Institute, MNI, and two from the *Centre de Recherche de l’Institut Universitaire de Gériatrie de Montréal*, CRIUGMa and CRIUMGb), and 2 samples with distinct acquisition protocols from the Alzheimer’s Disease Neuroimaging Initiative 2 (ADNI2a and ADNI2b) (Table 2). We selected subsamples of the MNI, CRIUGMa, CRIUGMb, ADNI2a and ADNI2b datasets such that patients and controls groups had identical sample size for each acquisition protocol or study, respectively 13, 13, 8, 20 and 11 subjects per group. The combined sample included 65 patients diagnosed with either amnestic MCI or AD dementia and 65 cognitively normal controls. Patients and controls were selected from a larger initial pool such that they would be matched for age, gender ratio as well as motion (see rs-fMRI preprocessing section). Distributions of age, gender and motion were as follows for patients vs. controls: age (mean ± std) = 72.7 ± 7.9 vs. 72.6 ± 7.3 years old, 41/24 vs. 41/24 females/males, residual frame displacement (mean ± std) = 0.22 ± 0.07 vs. 0.23 ± 0.08. All subjects gave informed consent to participate in these studies, which were approved by the research ethics committees of the institutions involved in data acquisition. Consent was obtained for data sharing and secondary analysis, the latter being approved by the ethics committee at the CRIUGM.

**Table 2.**
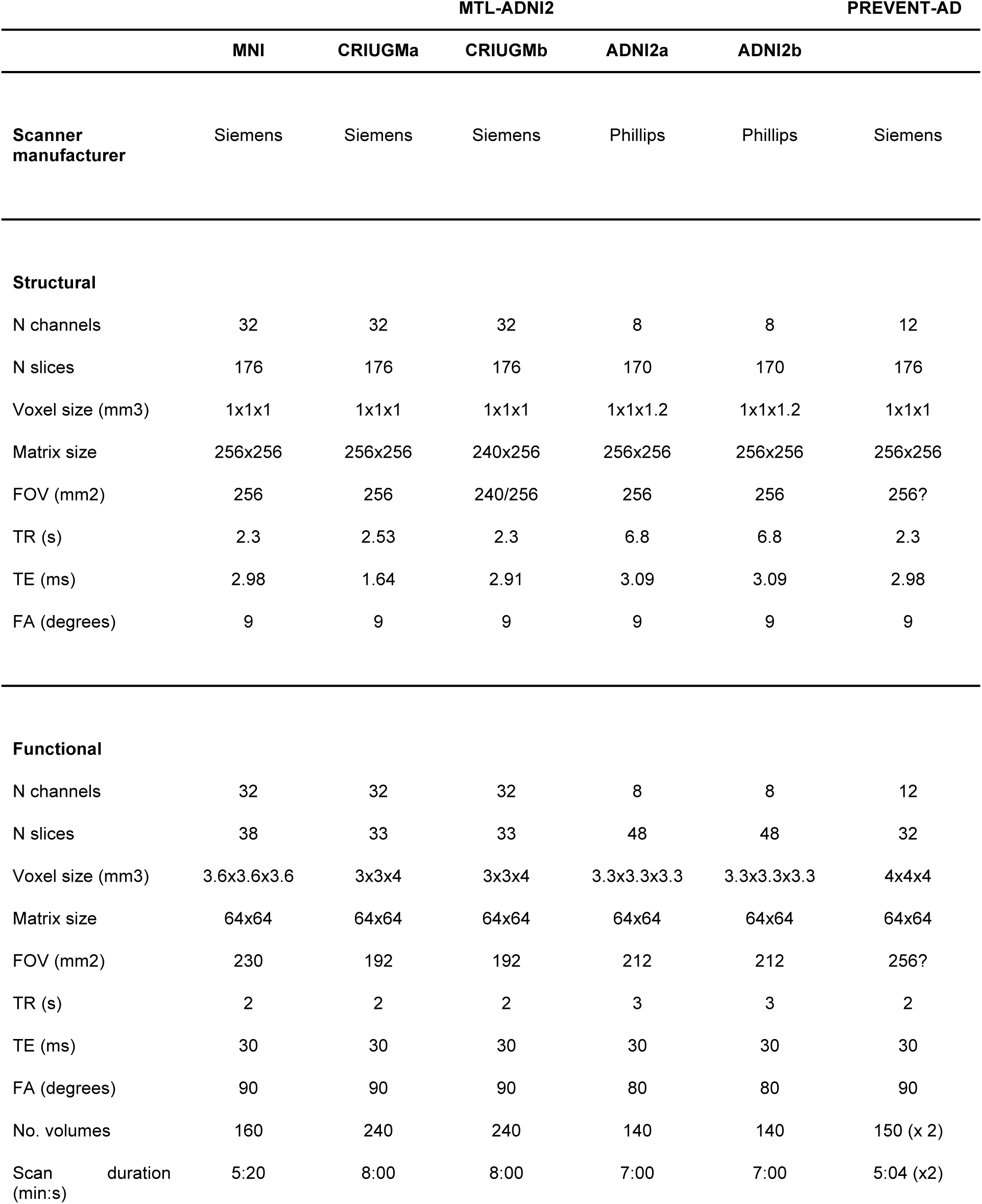
MRI acquisition protocols. Scan parameters are given for structural and functional data across the 5 MTL-ADNI samples as well as the PREVENT-AD dataset.

The PREVENT-AD dataset used in the present analysis included 231 cognitively healthy older adults with a known family history of AD, as reflected by a diagnosis of AD dementia in parent or first-degree relatives. PREVENT-AD participants were younger (mean ± std: 64.1 ± 5.7 years old) than subjects in the MTL-ADNI2 multisite sample and were not balanced for gender (172/59 females/males). All subjects had given informed consent and the study was approved by the "Research, Ethics and Compliance Committee" of McGill University.

### Note on the cohorts

The ADNI2 data used in the preparation of this article were obtained from the Alzheimer’s Disease Neuroimaging Initiative (ADNI) database (adni.loni.usc.edu). The ADNI was launched in 2003 by the National Institute on Aging (NIA), the National Institute of Biomedical Imaging and Bioengineering (NIBIB), the Food and Drug Administration (FDA), private pharmaceutical companies and non-profit organizations, as a $60 million, 5-year public-private partnership representing efforts of many co-investigators from a broad range of academic institutions and private corporations. A central goal of ADNI is to facilitate the discovery of biomarkers of very early AD progression, using MRI among other techniques. ADNI was followed by ADNI-GO and ADNI-2. In this study, we only included subjects from the two ADNI2 scanners (Achieva and Intera) associated with the largest samples. For up-to-date information, see www.adni-info.org.

The PREVENT-AD data were obtained from the Pre-symptomatic Evaluation of Novel or Experimental Treatments for Alzheimer’s Disease (PREVENT-AD) program data release 2.0 (November 30, 2015). The cohort of this program was composed of cognitively healthy individuals at increased risk of AD dementia because they have / had a first-degree relative (parent or sibling) who has / had dementia suggestive of AD. This cohort includes volunteers of age 60 or older (55 or older if current age is within 15 years of affected relative’s estimated age at onset of dementia). One current project consists of an observational study where participants are followed longitudinally once a year with a battery of tests and imaging modalities. In the present work, we focused on baseline data. A subset of test-retest rsfMRI data in 80 PREVENT-AD subjects has been shared publicly (Orban et al., 2015).

### Clinical evaluation

All subjects from the MTL-ADNI2 and PREVENT-AD samples underwent neuropsychological testing to assess cognitive function, including memory, language and executive abilities. However, the neuropsychological tests administered to participants varied across sites, as did criteria and clinical scales used for diagnosis of either MCI or AD. Briefly, patients with (amnestic) MCI had memory complaints and objective cognitive loss, yet showed intact functional abilities and did not meet criteria for a diagnosis of dementia in contrast with AD patients. HC demonstrated intact cognitive functions. Details on clinical evaluation for each cohort per site follow.

In ADNI2, the Mini-Mental State Evaluation (MMSE) and Clinical Dementia Rating (CDR) were used to distinguish between HC, MCI and AD subjects. MMSE scores were inclusively comprised between 24-30, 24-30 and 20-26 for HC, MCI and AD subjects, respectively. MCI patients had a CDR of 0.5 and AD patients a CDR of 0.5 or 1. An objective memory loss was evidenced with the Wechsler Memory Scale Logical Memory II in MCI, yet other cognitive domains and functional activities were unaffected. In addition, there was an absence of dementia, by contrast with AD patients who met the National Institute of Neurological and Communicative Disorders and Stroke / Alzheimer’s Disease and Related Disorders Association (NINCDS/ADRDA) criteria for probable AD (McKhann et al. 1984). The MNI sample only included MCI patients, who were similarly diagnosed using the MMSE, following Petersen Criteria (Petersen 2004). Subjects in the CRIUGM samples were administered the MMSE as well as the Montreal Cognitive Assessment (MoCA) (Nasreddine et al., 2005) and the Mattis Dementia Rating Scale (Schmidt et al. 1994). The diagnosis of MCI was made based on scores equal to or >1.5 standard deviations below the mean adjusted for age and education on memory tests, with input from a neurologist. A diagnosis of AD was determined according to the Diagnostic and Statistical Manual of Mental Disorders (4th ed.; American Psychiatric Association, 2000) and NINCDS/ADRDA clinical criteria, with input from a neurologist. Participants in the PREVENT-AD were evaluated for any cognitive impairment and symptoms suggestive of AD using the Repeatable Battery for the Assessment of Neuropsychological Status - RBANS (Randolph et al., 1998), the CDR, the MoCA and the AD8 Dementia screening (Galvin et al., 2005). Exclusion criteria common to all participants included contraindications to MRI, presence or history of neurologic disease with potential impact on cognition (e.g., Parkinson’s disease), and presence or history of substance abuse.

### Genetic and CSF biomarkers in PREVENT-AD subjects

In 228 PREVENT-AD subjects, DNA was isolated from 200 ul of whole blood using a QIASymphony apparatus and the DNA Blood Mini QIA Kit (Qiagen, Valencia, CA, USA). The standard QIASymphony isolation program was performed as per the manufacturer’s instructions. APOE single nucleotide polymorphism (SNP) genotyping was performed using pyrosequencing (PyroMArk96) and processed with GenomeStudio (version 2010.3) using standard methods.

CSF samples were obtained by lumbar puncture in 59 subjects of the PREVENT-AD cohort. For each subject, 25 ml of CSF was centrifuged 10 minutes +/- 2000g at room temperature and aliquoted in 50 vials of 0.5 ml and frozen at -80C for further analysis. Protein levels of Aβ_1-42_, total tau (tTau) and phosphorylated tau (pTau) were determined by enzyme-linked immunosorbent assay (ELISA) from Innotest technology (Fujirebio). These measurements were standardized with the European project BIOMARKAPD (Reijs et al., 2015), which intends to harmonize assays that are used to measure biological markers in neurodegenerative diseases.

### MRI acquisition

The MTL-ADNI2 multisite resting-state dataset included brain imaging data acquired on 3T MRI scanners (Table 2). Vendors differed between sites (Siemens Magnetom Tim Trio in MTL sites and Phillips Achieva or Intera in ADNI2). Analyses were performed on the first usable scan, typically the baseline scan when several scans were available. Functional scan acquisition parameters varied from one site to another, notably in run duration (ranges: 5min20s-8min), number of volumes (range: 140-240 vols), voxel size (range: 3-4x3-3.6x3.3-4mm^3^) and repetition time (range: 2-3s). Brain imaging data of the PREVENT-AD dataset were collected on a single 3T MRI scanner (Siemens, Magnetom Tim Trio). Two consecutive resting-state runs of 150 functional volumes were acquired, each run lasting 5min 04s. Spatial and temporal resolutions were as follows: voxel size = 4x4x4mm^3^ and repetition time = 2000ms. Table 2 reports scan acquisition parameters for all data.

### rs-fMRI preprocessing

Datasets were preprocessed and analyzed using the NeuroImaging Analysis Kit - NIAK - version 0.12.17 (http://niak.simexp-lab.org), under CentOS with Octave (http://gnu.octave.org) version 3.6.1 and the MINC toolkit (http://bic-mni.github.io/) version 0.3.18. Analyses were executed in parallel on the "Guillimin" supercomputer (http://www.calculquebec.ca/en/resources/compute-servers/guillimin), using the pipeline system for Octave and Matlab - PSOM (Bellec et al., 2012).

Each fMRI dataset was corrected for differences in timing of slice acquisitions; a rigid-body motion was then estimated using Minctracc (Collins and Evans, 1997) for each time frame, both within and between runs, as well as between one fMRI run and the T1 scan for each subject. The T1 scan was itself non-linearly co-registered to the Montreal Neurological Institute (MNI) ICBM152 stereotaxic symmetric template (Fonov et al., 2011), using the CIVET pipeline (Ad-Dab’bagh et al., 2006). The rigid-body, fMRI-to-T1 and T1-to-stereotaxic transformations were all combined to resample the fMRI in MNI space at a 3 mm isotropic resolution. To minimize artifacts due to excessive motion, all time frames showing an average frame displacement (FD) greater than 0.5 mm were removed (Power et al., 2012). The following nuisance covariates were regressed out from the fMRI time series: slow time drifts (basis of discrete cosines with a 0.01 Hz high-pass cut-off), average signals in conservative masks of the white matter and the lateral ventricles as well as the first principal components (accounting for 95% variance) of the six rigid-body motion parameters and their squares (Giove et al., 2009; Lund et al., 2006). The fMRI volumes were finally spatially smoothed with a 6 mm isotropic Gaussian blurring kernel. A more detailed description of the pipeline can be found on the NIAK website (http://niak.simexp-lab.org/pipe_preprocessing.html).

### Individual voxel-wise connectivity maps based on large-scale network templates

For all 361 subjects included in the analyses, we computed voxel-wise connectivity maps associated with each of 7 network templates extracted from a functional brain atlas generated on 200 healthy subjects (https://doi.org/10.6084/m9.figshare.1285615.v1). The atlas included cerebellar, limbic, visual, motor, default-mode, fronto-parietal and salience networks. For each subject and each network, a network connectivity map was obtained by computing the Fisher-transformed Pearson’s correlations between the average time course within the network template and the time course of every voxel in the brain grey matter. For each network, subject by voxel connectivity matrices were defined at the group level, separately for the MTL-ADNI and PREVENT-AD samples. Two general linear models were used to regress the following confounds on the group connectivity matrices: age, sex and residual (after scrubbing) FD, as well as acquisition protocols / study using dummy variables, i.e. MNI, CRIUGMa, CRIUGMb, ADNIa, ADNIb. The inclusion of constant terms in the models effectively normalized network connectivity maps to a zero grand mean across all subjects, separately for the MTL-ADNI and PREVENT-AD samples.

### Network subtypes defined by a cluster analysis in MTL-ADNI2 subjects

For each of the 7 rsfMRI networks, a subject by subject similarity (Pearson’s correlation) matrix summarized the between-subject correspondence of connectivity maps for all pairs of the 130 subjects in the MTL-ADNI multisite sample. A hierarchical cluster analysis was performed to identify 3 clusters of subjects whose network connectivity maps were similar in terms of spatial extent and/or strength. For each cluster, we defined a subtype of functional connectivity as the average connectivity map for subjects within this cluster. In total, there were thus 21 subtypes being investigated. Subtype weights were obtained by calculating the correlation between individual connectivity maps and each of the network subtype maps. Weights thus range between -1 and 1, with 1 meaning perfect correspondence, 0 lack of correspondence and -1 perfect but inverted correspondence.

### Statistical tests of association with clinical symptoms in MTL-ADNI2 subjects

We tested the association between subtypes of network connectivity and clinical symptoms in the 130 MTL-ADNI2 subjects. To this end, we employed two distinct statistical approaches: one approach treated subtypes as discrete units, where each subject belongs to one and only one cluster; a second approach used subtype weights, which are continuous measures. Despite these conceptual differences, we expected both statistical approaches to provide mostly concordant results. In the first approach, Chi2 tests were used to reveal unequal distributions of HC and ADMCI patients across the subtypes of each network. We report Cramer’s V effect sizes for which values of 0.1, 0.3 and 0.5 are respectively termed small, medium and large. In our second approach, we used general linear models to test separately the associations between the weights of each network subtype and clinical symptoms (HC vs. ADMCI). Because confounds (age, sex, rFD, sites) were regressed out prior to conducting this analysis, no factors of interest were entered in the general linear model. We provide Cohen’s f2 effect sizes for which values of 0.02, 0.15 and 0.35 are termed small, medium and large, respectively (Cohen, 1988). In both statistical approaches, results were deemed significant if they survived false-discovery rate (FDR) correction at q<0.05 across networks and subtypes.

### Matching of FH subjects to PAS subtypes

We next aimed to match connectivity maps in 231 cognitively normal FH older adults with PAS subtypes identified in the MTL-ADNI2 dataset. For each network and each PREVENT-AD subject, subtype weights were obtained by correlating his/her connectivity map (averaged over 2 runs) with each of the 3 subtype maps identified in the clinical sample. Each FH subject was assigned to the subtype for which the weight was maximal. We then tested, for each network, the similarity of subject distributions across subtypes between FH subjects in the PREVENT-AD cohort vs the distribution of ADMCI patients or HC subjects in the MTL-ADNI multisite sample. Chi2 tests were used to assess significance of differences in distributions and Cramer’s V values described effect sizes.

### Test-retest reliability of MTL-ADNI2 subtypes in FH subjects

Intra-class correlation coefficients quantified the reproducibility of weights between the two consecutive resting-state runs of the PREVENT-AD cohort. With 7 networks and 3 subtypes, we thus obtained 21 ICC measures. ICC measures were interpreted as follows (Cicchetti, 1994): less than 0.40 = poor, between 0.40 and 0.59 = fair, between .60 and 0.74 = good, between 0.75 and 1 = excellent.

### Statistical tests of association with AD biomarkers

We finally assessed whether the subtype weights of FH subjects would be associated with known biomarkers or risk factors of AD in PREVENT-AD. Namely, we investigated the possible association between APOE4 genotype, CSF Aβ_1-42_ and Tau levels with symptom associated network subtypes. Associations were tested in the framework of general linear models and were considered significant if they survived false-discovery rate (FDR) correction at *q*<0.05 across networks and subtypes. Because confounds (age, sex, rFD) were regressed out prior to conduct this analysis, no factors of interest were entered in the general linear models. Effect sizes are reported with Cohen’s f2 measures.

## Supplemental Information

NA

## Author contributions

Conceptualization, P.O., and P.B.; Data acquisition, M.S., C.M., A.S., A.D., J.P., P.R-N.; Analysis, P.O., A.T., M.S., C.M., C.D.; Writing - original draft, P.O., A.B., P.B.; Writing - review and editing, A.T., M.S., C.M., C.D., J.V., S.V., J.P.; Supervision, J.B., and P.B.; Funding, J.P., P.R-N., J.B., & P.B.

## Acknowledgments

Data collection and sharing for this project was funded by the Alzheimer’s Disease Neuroimaging Initiative (ADNI) (National Institutes of Health Grant U01 AG024904) and DOD ADNI (Department of Defense award number W81XWH-12-2-0012). ADNI is funded by the National Institute on Aging, the National Institute of Biomedical Imaging and Bioengineering, and through generous contributions from the following: Alzheimer’s Association; Alzheimer’s Drug Discovery Foundation; BioClinica, Inc.; Biogen Idec Inc.; Bristol-Myers Squibb Company; Eisai Inc.; Elan Pharmaceuticals, Inc.; Eli Lilly and Company; F. Hoffmann-La Roche Ltd and its affiliated company Genentech, Inc.; GE Healthcare; Innogenetics, N.V.; IXICO Ltd.; Janssen Alzheimer Immunotherapy Research & Development, LLC.; Johnson & Johnson Pharmaceutical Research & Development LLC.; Medpace, Inc.; Merck & Co., Inc.; Meso Scale Diagnostics, LLC.; NeuroRx Research; Novartis Pharmaceuticals Corporation; Pfizer Inc.; Piramal Imaging; Servier; Synarc Inc.; and Takeda Pharmaceutical Company. The Canadian Institutes of Health Research is providing funds to support ADNI clinical sites in Canada. Private sector contributions are facilitated by the Foundation for the National Institutes of Health (www.fnih.org). The grantee organization is the Northern California Institute for Research and Education, and the study is coordinated by the Alzheimer’s Disease Cooperative Study at the University of California, San Diego. ADNI data are disseminated by the Laboratory for Neuro Imaging at the University of Southern California. This research was also supported by NIH grants P30 AG010129 and K01 AG030514.

Data collection and sharing for the PREVENT-AD program were supported by its sponsors, McGill University, the Fonds de Research du Québec – Santé, the Douglas Hospital Research Centre and Foundation, the Government of Canada, the Canadian Foundation for Innovation, the Levesque Foundation, an unrestricted gift from Pfizer Canada Genome Quebec Innovation Center. Private sector contributions are facilitated by the Development Office of the McGill University Faculty of Medicine and by the Douglas Hospital Research Centre Foundation (http://www.douglas.qc.ca/).

This work was supported by a salary award (Junior 1 scholarship) by Fonds de Recherche du Québec - Santé (FRQS) as well as funds by the Canadian Institutes of Health research (CIHR), the Alzheimer’s Society of Canada and the Courtois foundation to PB. AB was supported by postdoctoral fellowships from the Canadian Alzheimer Society and CIHR, as well as by the Lemaire foundation. CD is supported by a bursary from the Lemaire foundation. We are grateful to Sylvie Belleville, Howard Chertkow, Louis Collins, Samir Das, Alain Dagher, Alan Evans, Alexandru Hanganu, Ouri Monchi, Amir Schmuel, Louise Theroux and Seqian Wang for sharing fMRI datasets and/or analytical tools.

Finally, we would like to acknowledge the participants of the PREVENT-AD cohort for dedicating their time and energy to help us collecting this data.

## References

Ad-Dab’bagh, Y., Lyttelton, O., Muehlboeck, J.S., Lepage, C., Einarson, D., Mok, K., Ivanov, O., Vincent, R.D., Lerch, J., Fombonne, E., Others, 2006. The CIVET image-processing environment: a fully automated comprehensive pipeline for anatomical neuroimaging research. In: Proceedings of the 12th Annual Meeting of the Organization for Human Brain Mapping. Florence, Italy, p. 2266.

Badhwar, A., Tam, A., Dansereau, C., Orban, P., Hoffstaedter, F., Bellec, P., 2017. Resting-state network dysfunction in Alzheimer’s disease: A systematic review and meta-analysis. Alzheimer’s & Dementia: Diagnosis, Assessment & Disease Monitoring 8, 73–85.

Badhwar, A., Tam, A., Dansereau, C., Orban, P., Toro, R., Bellec, P., 2016. Resting-state network dysfunction in Alzheimer’s disease: a systematic review and meta-analysis. Alzheimers. Dement.

Beach, T.G., Monsell, S.E., Phillips, L.E., Kukull, W., 2012. Accuracy of the clinical diagnosis of Alzheimer disease at National Institute on Aging Alzheimer Disease Centers, 2005-2010. J. Neuropathol. Exp. Neurol. 71, 266–273.

Bellec, P., Lavoie-Courchesne, S., Dickinson, P., Lerch, J.P., Zijdenbos, A.P., Evans, A.C., 2012. The pipeline system for Octave and Matlab (PSOM): a lightweight scripting framework and execution engine for scientific workflows. Front. Neuroinform. 6, 7.

Braak, H., Braak, E., 1991. Neuropathological stageing of Alzheimer-related changes. Acta Neuropathol. 82, 239–259.

Brier, M.R., Thomas, J.B., Ances, B.M., 2014. Network dysfunction in Alzheimer’s disease: refining the disconnection hypothesis. Brain Connect. 4, 299–311.

Buckner, R.L., Snyder, A.Z., Shannon, B.J., LaRossa, G., Sachs, R., Fotenos, A.F., Sheline, Y.I., Klunk, W.E., Mathis, C.A., Morris, J.C., Mintun, M.A., 2005. Molecular, structural, and functional characterization of Alzheimer’s disease: evidence for a relationship between default activity, amyloid, and memory. J. Neurosci. 25, 7709– 7717.

Chételat, G., La Joie, R., Villain, N., Perrotin, A., de La Sayette, V., Eustache, F., Vandenberghe, R., 2013. Amyloid imaging in cognitively normal individuals, at-risk populations and preclinical Alzheimer’s disease. Neuroimage Clin 2, 356–365.

Cicchetti, D.V., 1994. Guidelines, criteria, and rules of thumb for evaluating normed and standardized assessment instruments in psychology. Psychol. Assess. 6, 284.

Cohen, J., 1988. Statistical power analysis for the behavioral sciences Lawrence Earlbaum Associates. Hillsdale, NJ 20–26.

Collins, D.L., Evans, A.C., 1997. Animal: Validation and Applications of Nonlinear Registration-Based Segmentation. Int. J. Pattern Recognit Artif Intell. 11, 1271– 1294.

Dansereau, C., Benhajali, Y., Risterucci, C., Pich, E.M., Orban, P., Arnold, D., Bellec, P., 2017. Statistical power and prediction accuracy in multisite resting-state fMRI connectivity. Neuroimage 149, 220–232.

Delbeuck, X., Van der Linden, M., Collette, F., 2003. Alzheimer’disease as a disconnection syndrome? Neuropsychol. Rev. 13, 79–92.

Doan, N.T., Engvig, A., Persson, K., Alnæs, D., Kaufmann, T., Rokicki, J., Córdova-Palomera, A., Moberget, T., Brækhus, A., Barca, M.L., Engedal, K., Andreassen, O.A., Selbæk, G., Westlye, L.T., 2017. Dissociable diffusion MRI patterns of white matter microstructure and connectivity in Alzheimer’s disease spectrum. Sci. Rep. 7, 45131.

Dong, A., Toledo, J.B., Honnorat, N., Doshi, J., Varol, E., Sotiras, A., Wolk, D., Trojanowski, J.Q., Davatzikos, C., Alzheimer’s Disease Neuroimaging Initiative, 2017. Heterogeneity of neuroanatomical patterns in prodromal Alzheimer’s disease: links to cognition, progression and biomarkers. Brain 140, 735–747.

Dubois, B., Hampel, H., Feldman, H.H., Scheltens, P., Aisen, P., Andrieu, S., Bakardjian, H., Benali, H., Bertram, L., Blennow, K., Broich, K., Cavedo, E., Crutch, S., Dartigues, J.-F., Duyckaerts, C., Epelbaum, S., Frisoni, G.B., Gauthier, S., Genthon, R., Gouw, A.A., Habert, M.-O., Holtzman, D.M., Kivipelto, M., Lista, S., Molinuevo, J.-L., O’Bryant, S.E., Rabinovici, G.D., Rowe, C., Salloway, S., Schneider, L.S., Sperling, R., Teichmann, M., Carrillo, M.C., Cummings, J., Jack, C.R., Jr, Proceedings of the Meeting of the International Working Group (IWG) and the American Alzheimer’s Association on “The Preclinical State of AD”; July 23, 2015; Washington DC, USA, 2016. Preclinical Alzheimer’s disease: Definition, natural history, and diagnostic criteria. Alzheimers. Dement. 12, 292–323.

Elman, J.A., Madison, C.M., Baker, S.L., Vogel, J.W., Marks, S.M., Crowley, S., O’Neil, J.P., Jagust, W.J., 2016. Effects of Beta-Amyloid on Resting State Functional Connectivity Within and Between Networks Reflect Known Patterns of Regional Vulnerability. Cereb. Cortex 26, 695–707.

Filippini, N., MacIntosh, B.J., Hough, M.G., Goodwin, G.M., Frisoni, G.B., Smith, S.M., Matthews, P.M., Beckmann, C.F., Mackay, C.E., 2009. Distinct patterns of brain activity in young carriers of the APOE-ε4 allele. Proceedings of the National Academy of Sciences 106, 7209–7214.

Fonov, V., Evans, A.C., Botteron, K., Almli, C.R., McKinstry, R.C., Collins, D.L., Brain Development Cooperative Group, 2011. Unbiased average age-appropriate atlases for pediatric studies. Neuroimage 54, 313–327.

Galvin, J.E., Roe, C.M., Powlishta, K.K., Coats, M.A., Muich, S.J., Grant, E., Miller, J.P., Storandt, M., Morris, J.C., 2005. The AD8: a brief informant interview to detect dementia. Neurology 65, 559–564.

Giove, F., Gili, T., Iacovella, V., Macaluso, E., Maraviglia, B., 2009. Images-based suppression of unwanted global signals in resting-state functional connectivity studies. Magn. Reson. Imaging 27, 1058–1064.

Greicius, M.D., Srivastava, G., Reiss, A.L., Menon, V., 2004. Default-mode network activity distinguishes Alzheimer’s disease from healthy aging: evidence from functional MRI. Proc. Natl. Acad. Sci. U. S. A. 101, 4637–4642.

Hwang, J., Kim, C.M., Jeon, S., Lee, J.M., Hong, Y.J., Roh, J.H., Lee, J.-H., Koh, J.-Y., Na, D.L., Alzheimer’s Disease Neuroimaging Initiative, 2016. Prediction of Alzheimer’s disease pathophysiology based on cortical thickness patterns. Alzheimers. Dement. 2, 58–67.

Hyman, B.T., Phelps, C.H., Beach, T.G., Bigio, E.H., Cairns, N.J., Carrillo, M.C., Dickson, D.W., Duyckaerts, C., Frosch, M.P., Masliah, E., Mirra, S.S., Nelson, P.T., Schneider, J.A., Thal, D.R., Thies, B., Trojanowski, J.Q., Vinters, H.V., Montine, T.J., 2012. National Institute on Aging-Alzheimer’s Association guidelines for the neuropathologic assessment of Alzheimer’s disease. Alzheimers. Dement. 8, 1–13.

Jiang, Y., Huang, H., Abner, E., Broster, L.S., Jicha, G.A., Schmitt, F.A., Kryscio, R., Andersen, A., Powell, D., Van Eldik, L., Gold, B.T., Nelson, P.T., Smith, C., Ding, M., 2016. Alzheimer’s Biomarkers are Correlated with Brain Connectivity in Older Adults Differentially during Resting and Task States. Front. Aging Neurosci. 8, 15.

Jones, D.T., Knopman, D.S., Gunter, J.L., Graff-Radford, J., Vemuri, P., Boeve, B.F., Petersen, R.C., Weiner, M.W., Jack, C.R., Jr, Alzheimer’s Disease Neuroimaging Initiative, 2016. Cascading network failure across the Alzheimer’s disease spectrum. Brain 139, 547–562.

Korolev, I.O., Symonds, L.L., Bozoki, A.C., Alzheimer’s Disease Neuroimaging Initiative, 2016. Predicting Progression from Mild Cognitive Impairment to Alzheimer’s Dementia Using Clinical, MRI, and Plasma Biomarkers via Probabilistic Pattern Classification. PLoS One 11, e0138866.

Lam, B., Masellis, M., Freedman, M., Stuss, D.T., Black, S.E., 2013. Clinical, imaging, and pathological heterogeneity of the Alzheimer’s disease syndrome. Alzheimers. Res. Ther. 5, 1.

Lund, T.E., Madsen, K.H., Sidaros, K., Luo, W.-L., Nichols, T.E., 2006. Non-white noise in fMRI: does modelling have an impact? Neuroimage 29, 54–66.

Miller, K.L., Alfaro-Almagro, F., Bangerter, N.K., Thomas, D.L., Yacoub, E., Xu, J., Bartsch, A.J., Jbabdi, S., Sotiropoulos, S.N., Andersson, J.L.R., Griffanti, L., Douaud, G., Okell, T.W., Weale, P., Dragonu, I., Garratt, S., Hudson, S., Collins, R., Jenkinson, M., Matthews, P.M., Smith, S.M., 2016. Multimodal population brain imaging in the UK Biobank prospective epidemiological study. Nat. Neurosci. 19, 1523–1536.

Mufson, E.J., Malek-Ahmadi, M., Perez, S.E., Chen, K., 2016. Braak staging, plaque pathology, and APOE status in elderly persons without cognitive impairment. Neurobiol. Aging 37, 147–153.

Nasreddine, Z.S., Phillips, N.A., Bédirian, V., Charbonneau, S., Whitehead, V., Collin, I., Cummings, J.L., Chertkow, H., 2005. The Montreal Cognitive Assessment, MoCA: a brief screening tool for mild cognitive impairment. J. Am. Geriatr. Soc. 53, 695– 699.

Orban, P., Dansereau, C., Desbois, L., Mongeau-Pérusse, V., Giguère, C.-É., Nguyen, H., Mendrek, A., Stip, E., Bellec, P., 2017. Multisite generalizability of schizophrenia diagnosis classification based on functional brain connectivity. Schizophr. Res.

Orban, P., Madjar, C., Savard, M., Dansereau, C., Tam, A., Das, S., Evans, A.C., Rosa-Neto, P., Breitner, J.C.S., Bellec, P., PREVENT-AD Research Group, 2015. Test-retest resting-state fMRI in healthy elderly persons with a family history of Alzheimer’s disease. Sci Data 2, 150043.

Power, J.D., Barnes, K.A., Snyder, A.Z., Schlaggar, B.L., Petersen, S.E., 2012. Spurious but systematic correlations in functional connectivity MRI networks arise from subject motion. Neuroimage 59, 2142–2154.

Profenno, L.A., Porsteinsson, A.P., Faraone, S.V., 2010. Meta-analysis of Alzheimer’s disease risk with obesity, diabetes, and related disorders. Biol. Psychiatry 67, 505–512.

Randolph, C., Tierney, M.C., Mohr, E., Chase, T.N., 1998. The Repeatable Battery for the Assessment of Neuropsychological Status (RBANS): preliminary clinical validity. J. Clin. Exp. Neuropsychol. 20, 310–319.

Reijs, B.L.R., Teunissen, C.E., Goncharenko, N., Betsou, F., Blennow, K., Baldeiras, I., Brosseron, F., Cavedo, E., Fladby, T., Froelich, L., Gabryelewicz, T., Gurvit, H., Kapaki, E., Koson, P., Kulic, L., Lehmann, S., Lewczuk, P., Lleó, A., Maetzler, W., de Mendonça, A., Miller, A.-M., Molinuevo, J.L., Mollenhauer, B., Parnetti, L., Rot, U., Schneider, A., Simonsen, A.H., Tagliavini, F., Tsolaki, M., Verbeek, M.M., Verhey, F.R.J., Zboch, M., Winblad, B., Scheltens, P., Zetterberg, H., Visser, P.J., 2015. The Central Biobank and Virtual Biobank of BIOMARKAPD: A Resource for Studies on Neurodegenerative Diseases. Front. Neurol. 6, 216.

Scheltens, N.M.E., Galindo-Garre, F., Pijnenburg, Y.A.L., van der Vlies, A.E., Smits, L.L., Koene, T., Teunissen, C.E., Barkhof, F., Wattjes, M.P., Scheltens, P., van der Flier, W.M., 2016. The identification of cognitive subtypes in Alzheimer’s disease dementia using latent class analysis. J. Neurol. Neurosurg. Psychiatry 87, 235– 243.

Seeley, W.W., Crawford, R.K., Zhou, J., Miller, B.L., Greicius, M.D., 2009. Neurodegenerative diseases target large-scale human brain networks. Neuron 62, 42–52.

Selkoe, D.J., 2002. Alzheimer’s disease is a synaptic failure. Science 298, 789–791.

Sheline, Y.I., Morris, J.C., Snyder, A.Z., Price, J.L., Yan, Z., D’Angelo, G., Liu, C., Dixit, S., Benzinger, T., Fagan, A., Goate, A., Mintun, M.A., 2010. APOE4 allele disrupts resting state fMRI connectivity in the absence of amyloid plaques or decreased CSF Aβ42. J. Neurosci. 30, 17035–17040.

Sperling, R.A., Karlawish, J., Johnson, K.A., 2012. Preclinical Alzheimer disease—the challenges ahead. Nat. Rev. Neurol. 9, 54–58.

Stern, Y., 2006. Cognitive reserve and Alzheimer disease. Alzheimer Dis. Assoc. Disord. 20, S69–74.

Tampellini, D., 2015. Synaptic activity and Alzheimer’s disease: a critical update. Front. Neurosci. 9, 423.

Tanzi, R.E., 2012. The genetics of Alzheimer disease. Cold Spring Harb. Perspect. Med. 2.

Vemuri, P., Jones, D.T., Jack, C.R., Jr, 2012. Resting state functional MRI in Alzheimer’s Disease. Alzheimers. Res. Ther. 4, 2.

Wang, L., Brier, M.R., Snyder, A.Z., Thomas, J.B., Fagan, A.M., Xiong, C., Benzinger, T.L., Holtzman, D.M., Morris, J.C., Ances, B.M., 2013. Cerebrospinal fluid Aβ42, phosphorylated Tau181, and resting-state functional connectivity. JAMA Neurol. 70, 1242–1248.

Wang, L., Roe, C.M., Snyder, A.Z., Brier, M.R., Thomas, J.B., Xiong, C., Benzinger, T.L., Morris, J.C., Ances, B.M., 2012. Alzheimer disease family history impacts resting state functional connectivity. Ann. Neurol. 72, 571–577.

Yan, C.-G., Craddock, R.C., Zuo, X.-N., Zang, Y.-F., Milham, M.P., 2013. Standardizing the intrinsic brain: towards robust measurement of inter-individual variation in 1000 functional connectomes. Neuroimage 80, 246–262.

Zhang, X., Mormino, E.C., Sun, N., Sperling, R.A., Sabuncu, M.R., Yeo, B.T.T., Weiner, M.W., Aisen, P., Weiner, M., Petersen, R., Others, 2016. Bayesian model reveals latent atrophy factors with dissociable cognitive trajectories in Alzheimer’s disease. Proceedings of the National Academy of Sciences 113, E6535–E6544.

Bellec, P., Benhajali, Y., Carbonell, F., Dansereau, C., Albouy, G., Pelland, M., Craddock, C., Collignon, O., Doyon, J., Stip, E., Orban, P., 2015. Impact of the resolution of brain parcels on connectome-wide association studies in fMRI. Neuroimage 123, 212–228.

